# Facultative Cheating and Hybrid Vigor Resolves Cooperator-Cheater Conflict in a Yeast Public Goods System

**DOI:** 10.1101/2025.04.28.651155

**Authors:** Namratha Raj, Supreet Saini

## Abstract

The persistence of cooperation in the face of cheating is a central paradox in evolutionary biology. Microbial public goods systems employ diverse solutions to this dilemma, yet most studies assume fixed strategies wherein genotypes function strictly as cooperators or cheaters. Here, using the GAL/MEL regulon of *Saccharomyces cerevisiae*, we uncover a dynamic resolution to this conflict through facultative strategy switching. When haploid cheater-cooperator strains were co-evolved in melibiose, we observed the repeated emergence of same-mating-type diploid hybrids. These hybrids arise early in evolution and ultimately spread in the population. The hybrids exploit the public good produced by cooperator strains when present, acting as facultative cheaters. However, following cooperator extinction, hybrids switch to a cooperative phenotype. This dynamic role transition enables the hybrid to persist across shifting ecological contexts. Our findings reveal a novel, context-dependent mechanism of cooperation maintenance, whereby facultative cheating and genotype plasticity resolve the tension between individual fitness and collective benefit. This work expands the conceptual framework of social evolution by demonstrating that phenotypic flexibility, facilitated through hybridization, can stabilize cooperation even in fully exploitable public goods systems.

## Introduction

One of the most enduring puzzles in evolutionary biology is how cooperation is maintained in populations, particularly when it can be undermined by cheaters — individuals that reap the benefits of cooperation without paying its costs (1-4). In microbial populations, cooperative behaviors often manifest via secretion of enzymes or goods which serve a public function such as nutrient scavenging, degrading an antibiotic (5-8). Cells which contribute towards production and release of these public goods are referred to as cooperators, whereas those which simply benefit from the public goods released by other cells are referred to as cheaters.

Cooperation is thought to be evolutionarily unstable because cooperators are susceptible to exploitation by cheaters (9-11). Despite this inherent vulnerability, cooperative systems are widespread (3, 5, 12), suggesting that robust mechanisms must exist to sustain cooperation under natural conditions (13-18).

However, cheater-cooperator interactions have largely been framed in binary terms, where each genotype exhibits a fixed phenotype — either as a cooperator or as a cheater (19-25). This simplification, though analytically and experimentally convenient, overlooks the property of cells/populations to switch phenotypic commitment in response to environmental cues. The strategies by which genotypes can modulate cooperative investment in response to ecological or genetic context thus remains incompletely understood. Importantly, recent work has shown that other genotypes in the population can dramatically reshape the selective landscape, prompting shifts in metabolic investments, cooperative behavior, or competitive strategies (26, 27).

Public-goods driven extracellular hydrolysis of di- and tri-saccharides in yeast has been an important model system to study maintenance of cooperation (28-30). The *GAL/MEL* regulon in *Saccharomyces cerevisiae* is essential for growth in media where galactose and/or melibiose are the sole sources of carbon (31, 32). Melibiose — a disaccharide — is hydrolyzed extracellularly into glucose and galactose by α-galactosidase (Mel1p), encoded by the *MEL1* gene (**Figure 1A**). In well-mixed environments, the products of hydrolysis are equally available to all cells, creating a classic public goods scenario. Importantly, glucose represses, while galactose induces the expression of *MEL1*, setting up a regulatory landscape where metabolic feedback can influence cooperative behavior. Previous work has shown that such an environment drives metabolic diversification among members of an isogenic population, followed by genetic diversification (33).

**Figure 1.**
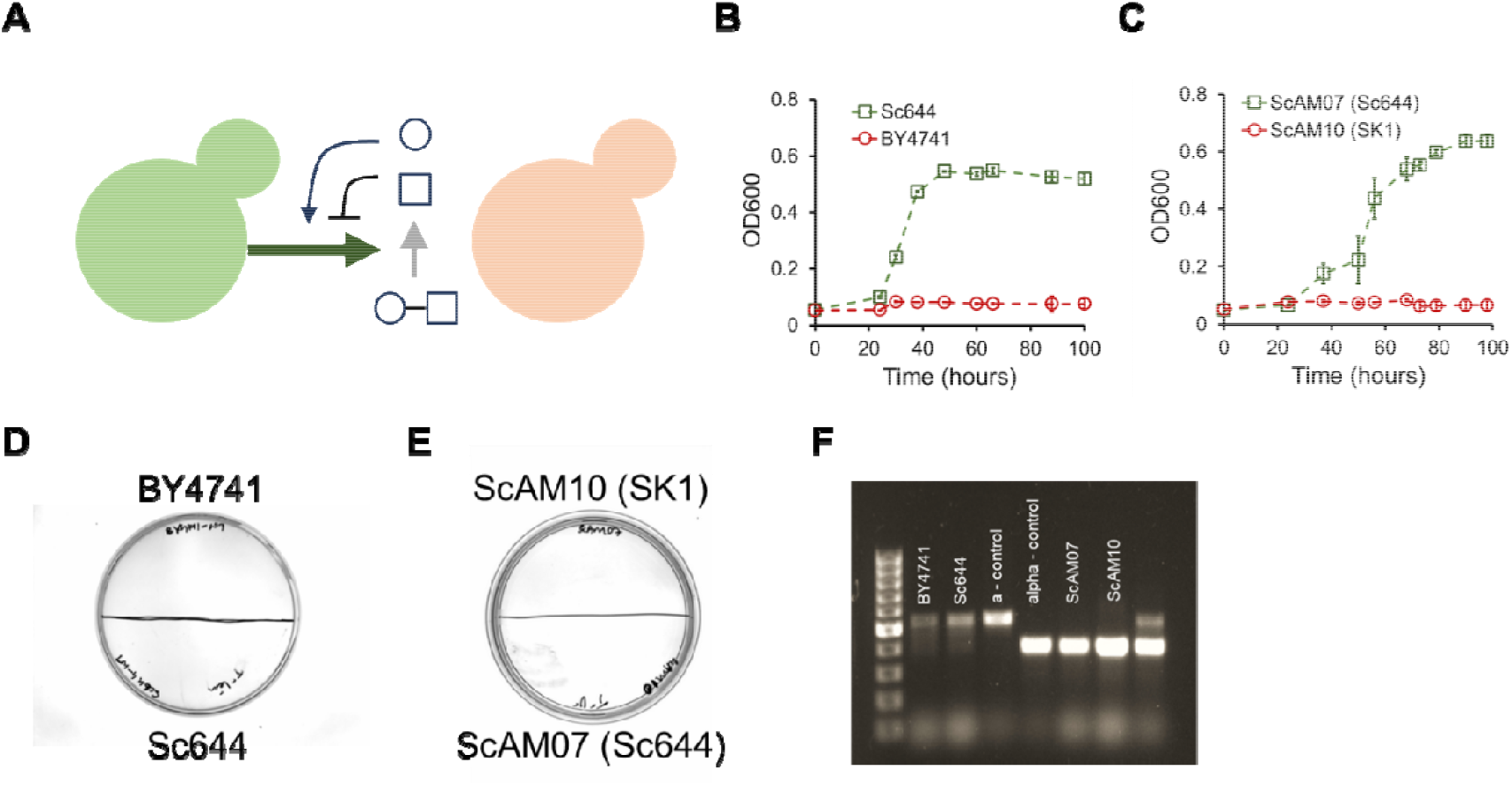
(A) Schematic representation of interactions between cooperator and cheater yeast cells in an environment containing melibiose as the carbon source. The cooperator cell expresses the *MEL1* gene and secretes the enzyme α-galactosidase (Mel1p), which hydrolyzes extracellular melibiose into its monosaccharide constituents glucose and galactose. These monosaccharides are then available for uptake by both cooperator and non-producing cheater cells. While the cooperator bears the metabolic cost of public goods production, the cheater utilizes the released sugars without incurring such costs. The regulatory network governing *MEL1* expression is influenced by the monosaccharides glucose and galactose. Glucose acts as a repressor, downregulating *MEL1* transcription, whereas galactose functions as an inducer, upregulating *MEL1*. This feedback creates a dynamic regulatory loop, whereby the products of melibiose hydrolysis simultaneously modulate cooperative behavior, potentially shaping population-level dynamics and the evolution of cooperative strategies. **(B, C)** The cheater and cooperator strains of yeast were grown in 2% melibiose. Growth assay confirms that cheater cannot grow in melibiose while cooperator grows independently. Growth kinetics of the pair BY4741 and Sc644-WT (B) and strains ScAM10 and ScAM07 (C). **(D, E)** The haploid strains used in this study do not grow in the amino acid double drop out plates both the strains are auxotrph for. **(F)** *MAT* locus of the haploid strains used in the co-evolution was screened. The first pair BY4741 and Sc644-WT are of ‘***a****’* mating type, while the second pair ScAM10 and ScAM07 are of ‘α’ mating type.

In the current study, we use the *GAL/MEL* system to explore the evolution of cooperation through a co-evolution experiment between a haploid *MEL1*+ cooperator strain and haploid *MEL1-* cheater strains of *S. cerevisiae*. After 200 generations of co-culture in melibiose, we observed the emergence of a novel genotype: hybrid diploid formed through mating after a *HO*-independent mating-type switch (34). The resulting hybrid is fitter in environments containing melibiose, thus outperforming both parental haploid strains. Our results show that these hybrids behaved as facultative cheaters—initially exploiting the public goods produced by cooperator strains. However, once the cooperators were driven to extinction, the hybrids themselves began producing the public good, effectively switching roles to become cooperators.

Our work reveals a novel, dynamic mechanism by which cooperation can be maintained: a genotype flexibly shifts between cheating and cooperating depending on the population context. Such facultative transitions blur the binary distinction between cheaters and cooperators and highlight the importance of genotype-by-environment interactions in shaping evolutionary outcomes. Our findings add a new dimension to the understanding of microbial cooperation and suggest that hybridization may serve not only as a source of genetic diversity but also as a mechanism for adaptive behavioral plasticity.

## Results

### Cooperation is Essential for Growth on Melibiose in *MEL1*-Deficient Yeast Strains

To investigate the dynamics of cooperation and cheating in yeast, we employed a system based on melibiose utilization, where only strains possessing the *MEL1* gene can secrete α-galactosidase (Mel1p) to hydrolyze melibiose into glucose and galactose. We used *Sc644* and its derivative *ScAM07* — both of which contain a functional *MEL1* gene — as representative cooperator strains. In contrast, we selected two commonly used laboratory strains, *ScAM10* (a derivative of SK1) and *BY4741* (S288c background), both of which lack *MEL1* and thus serve as cheaters.

To determine the growth capabilities of these strains in melibiose, we performed growth assays in a defined medium containing 2% melibiose as the sole carbon source. The results clearly indicated that *MEL1*-deficient strains were incapable of growth in melibiose, while *Sc644* and *ScAM07* showed robust growth, confirming their role as cooperators that produce the necessary public good (**Figure 1B** and **1C**).

To facilitate tracking of individual strains in co-cultures, we employed complementary auxotrophic markers in our cooperator-cheater pairs. For instance, *Sc644* is auxotrophic for tryptophan, and *BY4741* is auxotrophic for leucine. Consequently, neither strain can grow on media lacking both tryptophan and leucine, allowing us to unambiguously detect hybrid formation through colony growth on double-dropout plates. The inability of either haploid to grow on such media was verified through independent streaking experiments (**Figure 1D** and **1E**). Additionally, we verified the mating types of each strain pair via PCR to ensure both members of each pair were of the same mating type (either **a** or α), a critical control since same-mating-type fusion is rare. All strains were confirmed to be haploid using flow cytometry, establishing a controlled baseline for co-evolution experiments (**Figure 1F**).

### Emergence of Diploid Hybrids in Melibiose Co-evolution Cultures

Co-evolution experiments were initiated in melibiose medium with a 1:1 mixture of cooperator and cheater strains, and the cultures were propagated for 200 generations. At regular intervals, we sampled the populations to assess the frequency of each strain and to identify any emerging genotypes. After just one round of growth (∼6–7 generations), we observed the appearance of colonies on double-dropout media, suggesting hybrid formation. These colonies, which were otherwise absent when either haploid strain was plated alone, began to appear in all replicate lines, regardless of the specific strain pair used or their mating type (**Figure 2A**). Typically, 1 to 10 such colonies were observed among approximately 10□ cells per plate.

**Figure 2.**
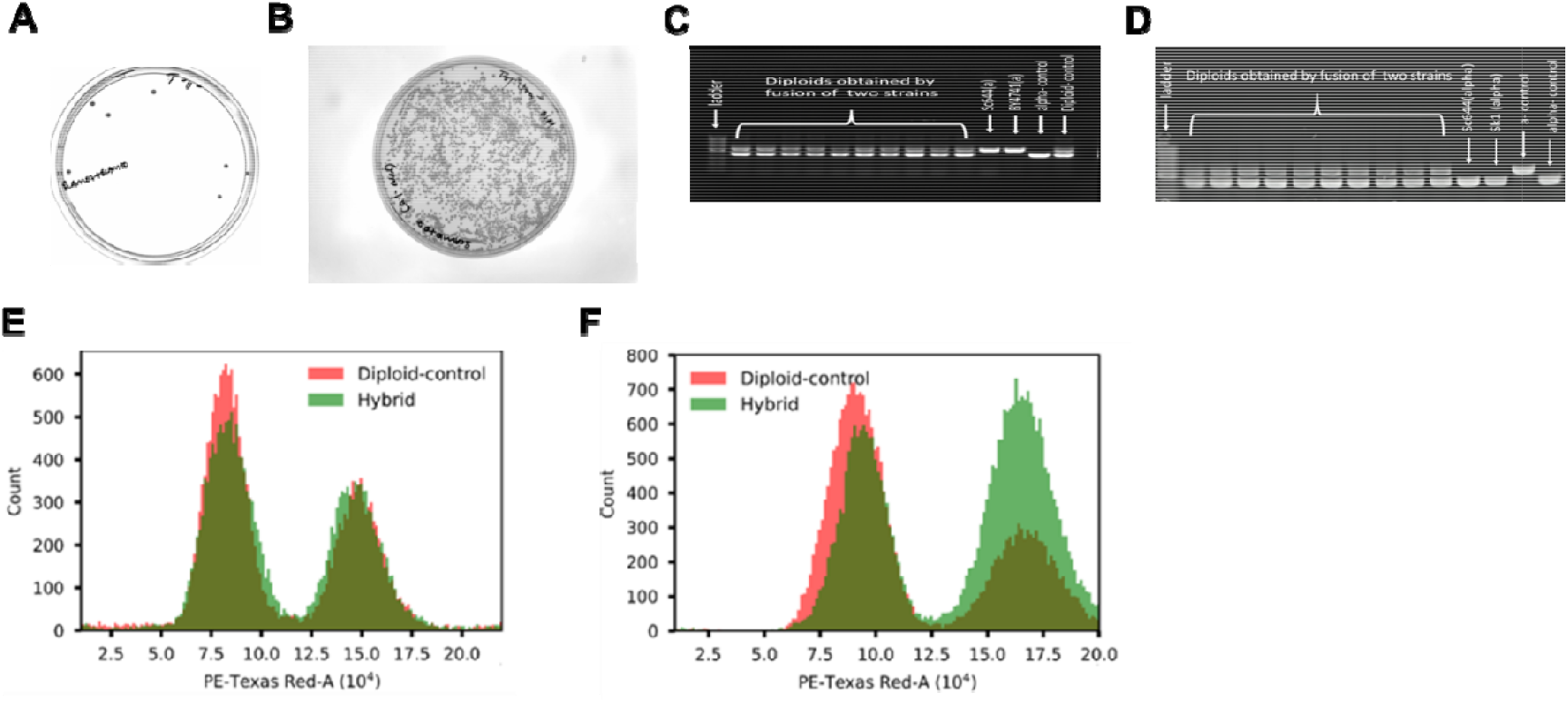
Validation of yeast hybrid generation through morphological, molecular, and cytometric analyses. **(A–B)** Visual confirmation of hybrid colony formation. Colonies on double dropout media confirm presence of diploid soon after the start of the experiment (A), and fixation of the hybrid after 200 generations (B). No haploid was detected in the media at 200 generations. **(C–D)** PCR-based genotypic validation of hybrid status. Gel electrophoresis shows hybrid strains carrying both MATa and MATα. **(E–F)** Flow cytometry analysis of DNA content in hybrids (green) compared to diploid controls (red). PE-Texas Red-A signal intensity, proportional to DNA content confirms diploidization.

The frequency of hybrid colonies increased progressively with each transfer. By the 200^th^ generation, hybrids constituted the majority of the population in every replicate, suggesting a strong selective advantage or persistent formation under the conditions of the experiment (**Figure 2B**).

To confirm that these colonies were indeed hybrids and not revertants that had lost auxotrophy, we isolated 10–20 colonies from each co-evolution line and analyzed their mating-type loci via PCR. For the *Sc644* and *BY4741* pair, the hybrids were predominantly heterozygous (a/α), indicative of a diploid. However, in co-cultures of *ScAM07* and *ScAM10* — both α mating type — we observed a more complex pattern. By generation 50, several lines contained homozygous α/α hybrids, while others exhibited heterozygous MAT loci, suggesting additional rounds of fusion or potential mating-type switching (**Figure 2C** and **2D**).

To further verify the diploid status of these hybrids, we performed flow cytometry-based ploidy analysis. All hybrids displayed fluorescence intensities consistent with diploid controls, clearly distinguishing them from haploid progenitors (**Figure 2D**). These findings demonstrate that same-mating-type fusions, although rare, can occur and produce stable diploid hybrids under selective pressure, such as dependency on public goods.

### Hybrid Formation Occurs Independently of Cooperative Dynamics

To determine whether hybridization was specifically driven by the cooperative interactions necessitated by melibiose metabolism, we conducted similar control co-evolution experiments in media containing 2% glucose and 2% galactose — i.e., carbon sources that both strains can metabolize independently. These media do not require public goods for growth, thereby eliminating the selective pressure for cooperative interactions. Surprisingly, hybrid colonies also emerged in these conditions, with frequencies and dynamics of frequency change similar to those observed in melibiose. This demonstrates that hybrid formation is not contingent on public goods-based cooperation but is instead an inherent potential outcome of yeast co-culture over time, possibly driven by rare mating-type switching.

Previous studies have described low-frequency, *HO*-independent mating-type switching events occurring at rates of approximately 1 in 10□ cells (35, 36). In our system, with total cell numbers approaching 10□ per co-culture, such events may be sufficient to enable cell fusion between strains of originally identical mating type. The repeated emergence and rise of diploid hybrids across all sugar environments supports the view that this phenomenon is both robust and generalizable across different genetic backgrounds and environmental conditions.

### Hybrid Vigor and Facultative Cheating Enable Hybrids to Outcompete Parental Haploids

To investigate why hybrids progressively outcompeted their haploid progenitors, we assessed their growth kinetics in all three sugar environments (glucose, galactose, and melibiose). In both glucose and galactose, hybrids demonstrated significantly higher growth rates and greater biomass accumulation compared to the two haploid strains, indicating hybrid vigor or heterosis (**Figure 3A-F**). However, in melibiose, hybrids displayed a notably prolonged lag phase — more than twice that of the haploid cooperator — suggesting a reduced capacity to initiate growth independently in the absence of pre-hydrolyzed sugars.

**Figure 3.**
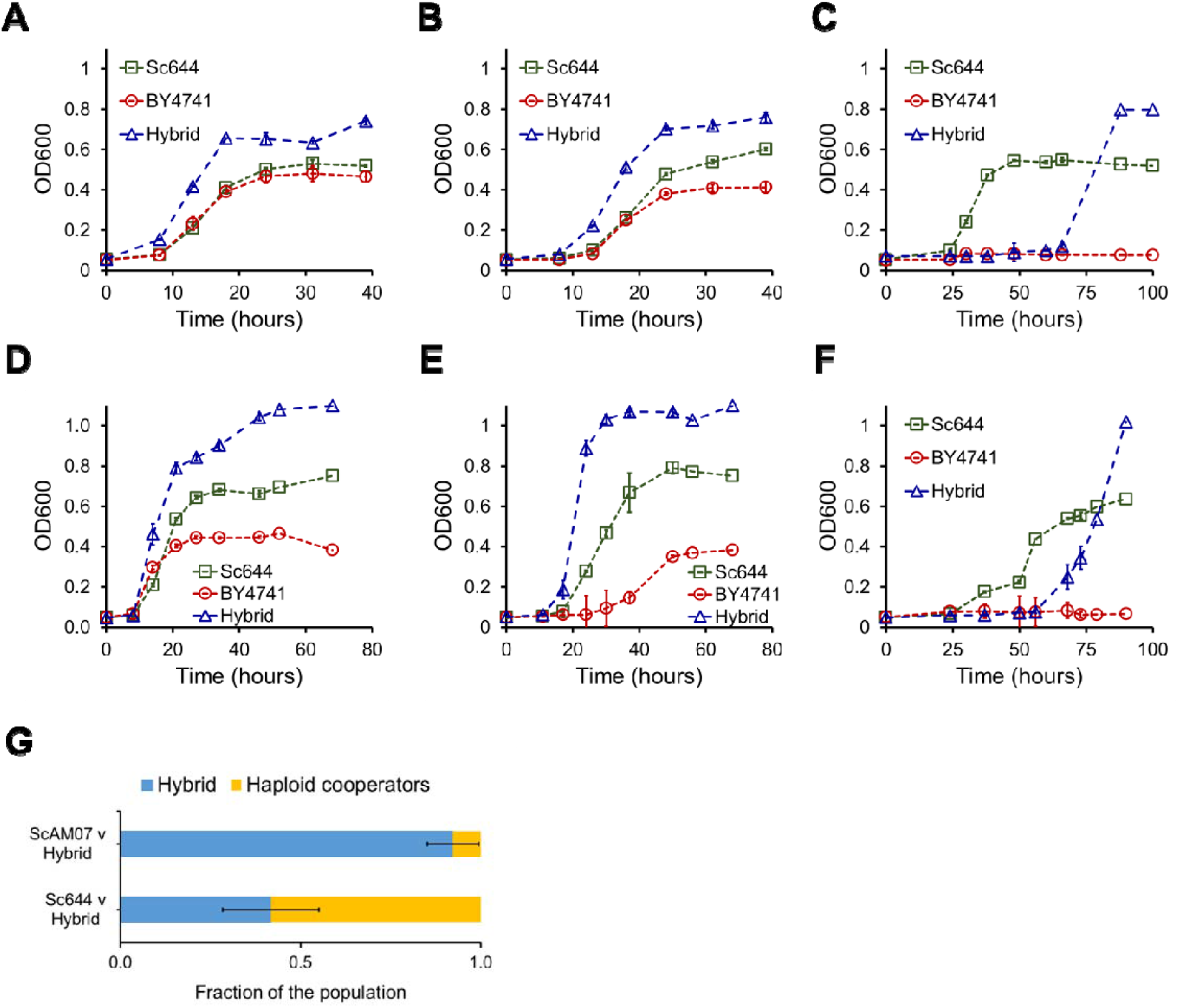
Comparative growth dynamics and population composition of yeast hybrids and parental haploids under different nutrient environments. **(A–F)** Growth kinetics of *Saccharomyces cerevisiae* hybrid strains (blue triangles) compared to their haploid parental strains Sc644 (cooperator, indicated by green squares) and BY4741 (cheater, indicated by red circles). (A-C) represent growth kinetics in (A) glucose; (B) galactose; and (C) melibiose for ScAM07 cooperator and BY4741 cheater. **(D-F)** represent growth kinetics in (D) glucose; (E) galactose; and (F) melibiose for ScAM07 cooperator and ScAM10 cheater. **(G)** Bar graphs quantify population compositions from competition experiments between hybrid and haploid strains. Two independent hybrid strains (ScAM07 and Sc644-derived hybrids) were co-cultured with haploid cooperators under nutrient-limited conditions. Population fractions were determined at the end of the experiment, revealing a strong dominance of hybrid cells (blue) over haploid cooperators (yellow), suggesting enhanced competitive fitness of hybrids.

Despite this fitness cost in melibiose, hybrids rapidly gained frequency when co-cultured with *MEL1+* haploid cooperators. To explore this further, we co-inoculated glycerol-lactate pre-grown hybrids and cooperators at a 1:1 ratio and monitored the frequency of hybrids after ∼30 hours of growth, corresponding to mid-exponential phase. We observed that hybrids had initiated growth and constituted a significant fraction of the population (**Figure 3G**). This is despite the hybrids exhibiting a lag phase of up to 60 hours, when grown by themselves. In contrast, cooperators entered exponential growth at around 25 hours.

These results strongly suggest that hybrids initially exploit the monosaccharides released by the cooperator through Mel1p activity, acting as facultative cheaters. Importantly, since hybrids retain the *MEL1* gene, they retain the latent ability to cooperate, potentially shifting their strategy based on population composition and resource availability.

In summary, our results reveal a previously uncharacterized mode of adaptation in microbial communities: hybrids formed through rare same-mating-type fusions can act as facultative cooperators. Their hybrid vigor allows them to outcompete parental strains in environments where public goods are available, even when their direct contribution to public goods production is delayed or minimal. These findings have broad implications for understanding microbial evolution in spatially and temporally heterogeneous environments where cooperation, cheating, and hybridization intersect.

## Discussion

Our study reveals a novel mechanism for the maintenance of cooperation in microbial populations, wherein genotypic plasticity facilitates dynamic transitions between cheating and cooperation. Using the *GAL/MEL* regulon of *Saccharomyces cerevisiae*, we demonstrate that co-evolving cooperator and cheater strains in a public goods system centred on melibiose leads to the spontaneous emergence of diploid hybrids via rare same-mating-type fusion. These hybrids initially act as facultative cheaters, exploiting the extracellular α-galactosidase activity of *MEL1+* cooperators to grow on hydrolyzed monosaccharides. However, upon cooperator extinction, the hybrids express *MEL1* themselves, adopting the cooperative role. This facultative shift allows hybrids to first capitalize on public goods and later ensure their own survival through cooperation, ultimately leading to their fixation in the population and the loss of both original haploid lineages.

These findings have broad implications for our understanding of the evolutionary dynamics underpinning cooperation (37, 38). The assumption that microbial genotypes exhibit fixed behavioral strategies — either cheater or cooperator — has shaped much of the theoretical and empirical literature (although context-dependent cooperative behavior has been studied in social interactions (39, 40)). Our results challenge this binary framing by uncovering a mechanism where a single genotype alters its cooperative phenotype based on ecological context. Such plasticity could be widespread in nature, especially in environments with fluctuating public goods availability, and may contribute to the resilience and stability of microbial communities (41-44).

In our work, adaptation happened because of hybridization. Autodiploidization of the cheater or the cooperator were not adaptive in melibiose. The diploid of the ancestor Sc644 exhibited identical growth rates and biomass as the haploid ancestor (p-value = 0.49; Welch’s t-test). However, hybrid from our experiment exhibited a longer lag and a higher biomass, as compared to the ancestor (p-value < 0.01, Welch’s t-test). The hybrid not only grew robustly but also exhibited higher biomass, as compared to the ancestor cooperator (45). Hybrid vigor has been known to dictate evolutionary routes in yeast populations in a variety of contexts (45-47).

The ability of hybrids to exhibit greater fitness in melibiose, glucose, and galactose further underscores the role of metabolic plasticity in shaping evolutionary outcomes. However, adaptation does not necessarily require hybrids. In several cases, simply a transition from haploid to diploid is adaptive (48-50). Although cases where a haploid is adaptive over a diploid are also known (51). This metabolic reprogramming in the hybrid during the course of growth with the cooperator is not only relevant to cooperative dynamics but also across broader biological themes such as, phenotypic plasticity in metabolism underlies critical processes such as adaptation to nutrient fluctuations, antibiotic resistance, and host-pathogen interactions across diverse microbial and multicellular systems (33, 52-56).

While we observe consistent hybrid formation and phenotypic shifts across multiple replicates and environments, the molecular basis for same-mating-type fusion and subsequent mating-type switching are not clearly understood. *HO*-independent mating-type switch in a haploid, leading to diploids, is known to occur with a small probability (34, 35). As a result, yeast geneticists have historically relied on using *STE3*Δ to prevent same-sex mating in a haploid population (57-59). While our experimental setup is constrained to well-mixed, nutrient-defined laboratory conditions, natural environments, with their spatial structure, ecological complexity, and interactions with additional microbial taxa may influence the dynamics of hybridization and cooperation in other more complex ways. Finally, while our classification of the hybrid’s behavior as “facultative cheating” is inferred from growth assays and population shifts; single-cell analyses of gene expression or public goods production would provide more details and the metabolic commitment of the population at a finer resolution.

In sum, our findings reveal a context-dependent evolutionary strategy that bridges cheating and cooperation through hybridization and metabolic plasticity. This mechanism expands the repertoire of solutions to the evolutionary dilemma of cooperation and suggests that the interplay between genetic fusion and environmental feedback can generate emergent, adaptable phenotypes that reshape social dynamics in microbial populations. Future work should explore whether similar facultative strategies emerge in other public goods systems and how such strategies affect long-term community structure and evolutionary stability.

## Material and Methods

### Strains Used

The experiments utilized *Saccharomyces cerevisiae* strains, derived from three genetic backgrounds: SK1, Sc644, and BY4741. The SK1-derived strain used was *ScAM10*, while *ScAM07* was derived from *Sc644*, a hybrid strain formed by mating *S. cerevisiae* with *S. carlsbergensis* (60). The third strain, *BY4741*, is a commonly used S288c-derived laboratory strain (61). A detailed list of strain genotypes and auxotrophic markers is provided in **Table 1**.

**Table 1.** Strains used in the study.

| **S. No.** | **Strain** | **Parent strain** | **MAT locus** | **Marker (+)** | **Marker (-)** |
| --- | --- | --- | --- | --- | --- |
| 1 | ScAM07 | Sc644 | $\alpha$ | *URA3* | *TRP1* |
| 2 | ScAM10 | SK1 | $\alpha$ | *TRP1* | *URA3* |
| 3 | BY4741 | S288c | a | *TRP1* | *LEU2* |
| 4 | Sc644-WT | Sc644 | a | *LEU2* | *TRP1* |

**Table 2.**
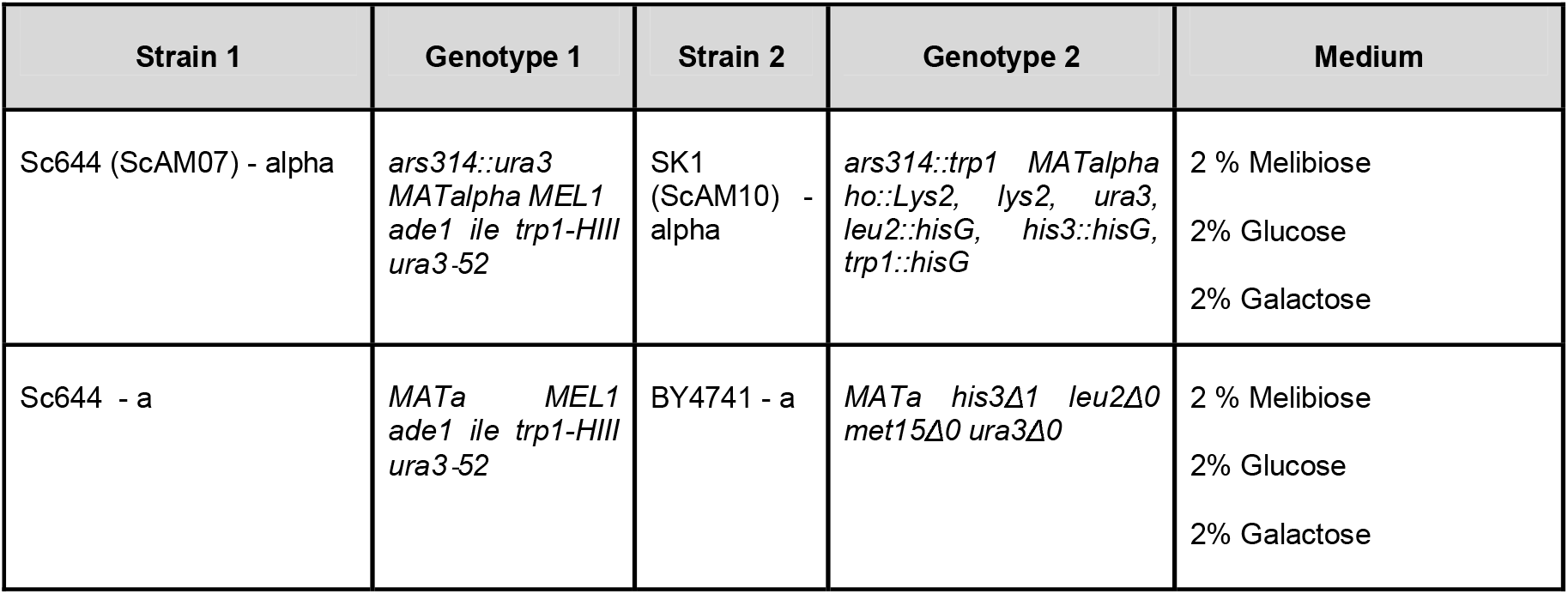
The genotype of the pairs of strains used in co-evolution study in various growth medium.

**Table 3.**
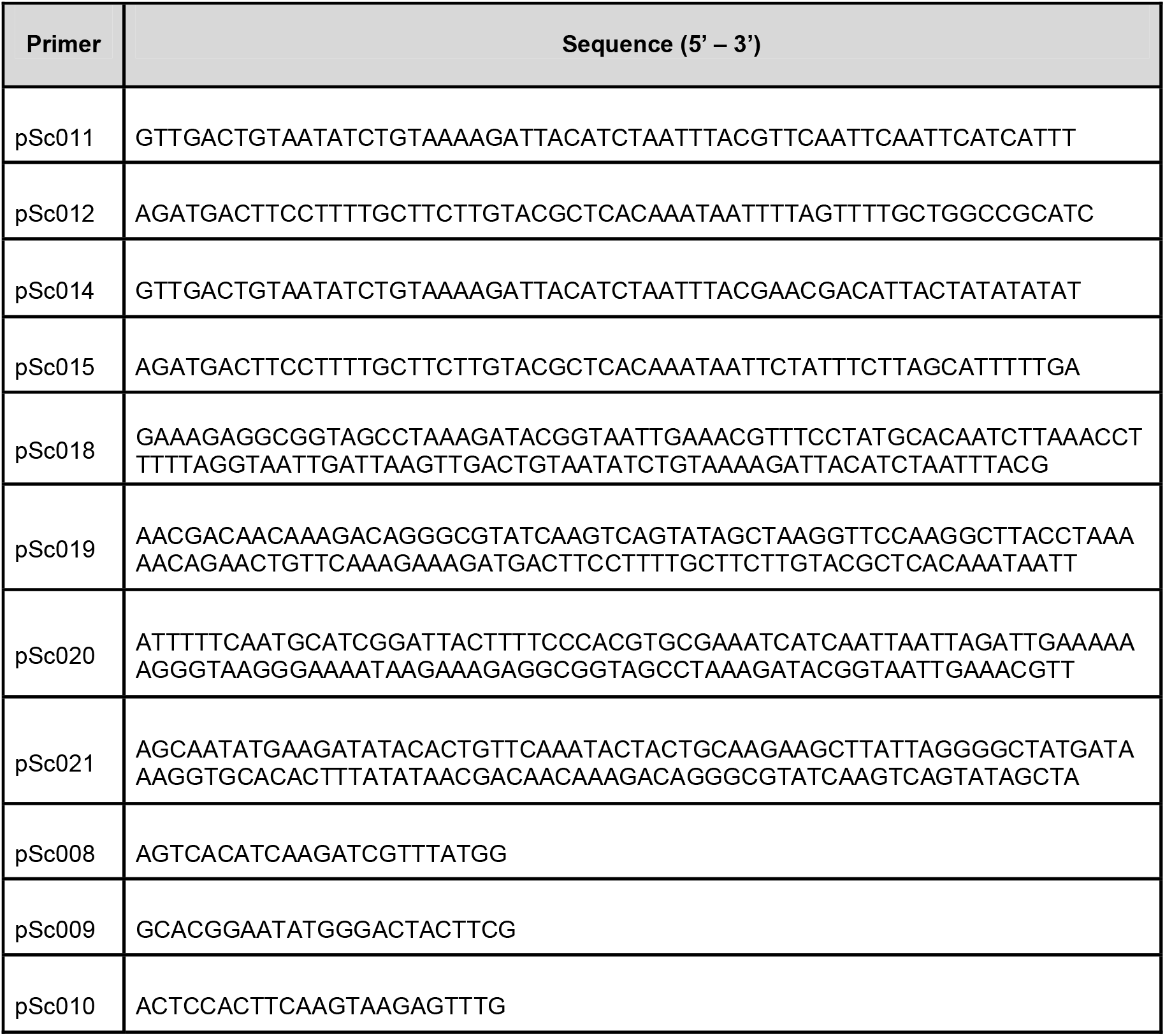
List of primers used to construct various strains used in the co-evolution study.

*ScAM07* was constructed by inserting a *URA3* marker into *Sc644*, which is auxotrophic for tryptophan and uracil, as previously described (49). *ScAM10*, derived from SK1, carries a *TRP1* marker and is auxotrophic for tryptophan, leucine, and uracil. The *URA3* and *TRP1* markers were inserted at the ARS314 locus on chromosome III near the MAT locus (62). For marker insertion, *URA3* was amplified from plasmid p426GPD (63) using primers pSc011 and pSc012, while *TRP1* was amplified from plasmid p424TEF (64) using primers pSc014 and pSc015. To ensure proper homologous recombination, extended flanking sequences were generated using primer pairs pSc018/pSc019 and subsequently pSc020/pSc021. PCR products were introduced into the relevant haploid strains by electroporation using an Eppendorf Electroporator.

### Co-evolution Experiment

Each co-evolution experiment began by reviving the yeast strains from frozen stocks on YPD agar (0.5% yeast extract, 1% peptone, and 2% dextrose). After 48 hours of growth, single colonies were inoculated into non-inducing, non-repressing glycerol-lactate medium and incubated for 42–48 hours. Co-evolution was then carried out in three sugar environments— 2% melibiose, 2% glucose, or 2% galactose—using complete synthetic media (CSM: 0.671% yeast nitrogen base with ammonium sulfate, 0.05% amino acid mixture). Equal volumes (50 µL each) of pre-grown cooperator and cheater cultures were mixed in a 1:1 ratio and inoculated into 5 mL of CSM containing one of the sugars. Five independent replicate co-evolution lines were established per sugar condition.

Cultures were incubated at 30°C with shaking at 250 rpm for 24 hours, after which they were propagated by transferring 1% (1:100 dilution) into fresh medium. Each such dilution corresponded to approximately 6.7 generations. This process was continued for 200 generations, with aliquots of each co-culture frozen every 50 generations for later analysis.

### Detection of Hybrids and MAT Locus Screening

To detect hybrid formation in the co-cultures, cells were plated on amino acid double-dropout plates tailored to the specific auxotrophies of each strain pair. Only hybrids, which have regained prototrophy through genetic complementation, can grow on such plates, while the original haploids cannot. Colonies observed on these plates were presumed hybrids. To confirm their genotype, ∼10 colonies from each co-evolution line were selected and subjected to PCR-based screening of the MAT locus. Primers used were pSc008 (flanking MAT locus), pSc009 (specific to MATα), and pSc010 (specific to MATa).

### Ploidy Analysis via Flow Cytometry

To determine the ploidy of a strain, colonies were cultured in YPD medium. After reaching an OD600 of ∼1.0 (measured with an Eppendorf Nano-spectrophotometer), 1.5 mL of culture was harvested, washed, and resuspended in 100 µL sterile water. Subsequently, 1 mL of 70% ethanol was added gradually while vortexing, and the suspension was incubated overnight at 4°C to fix the cells.

The next day, cells were washed with 500 µL RNase buffer (0.2 M Tris-Cl, 20 mM EDTA, pH 8.0) and resuspended in 100 µL of the same buffer. RNase A was added to a final concentration of 1 µg/µL, and samples were incubated at 37°C for 4 hours. Following RNase treatment, cells were washed with PBS and resuspended in 950 µL buffer. For DNA staining, 50 µL of 1 mg/mL propidium iodide was added to achieve a final concentration of 50 µg/mL. After 30 minutes of incubation at room temperature, the suspension was vortexed and sonicated briefly. Samples were then analyzed on a BD FACSAria SORP flow cytometer to assess DNA content and ploidy.

### Growth Assays

To evaluate growth dynamics, strains were revived on YPD plates and inoculated into glycerol-lactate medium, where they were grown to saturation (approximately 48 hours). These cultures were then diluted 1:100 into the desired test medium (melibiose, glucose, or galactose) to a starting OD600 of 0.05. Growth was monitored every 6–8 hours by measuring optical density at 600 nm using a Thermo Scientific Multiskan GO microplate reader until cultures reached saturation. These assays were used to compare growth kinetics among cooperator, cheater, and hybrid strains under different nutrient conditions.

### Competition Assays

To quantify relative fitness in mixed populations, cooperator and cheater strains were revived and pre-cultured in glycerol-lactate medium for 42–48 hours. Equal volumes (50 µL) of each culture were mixed in a 1:1 ratio and inoculated into 5 mL of CSM containing either 2% melibiose, glucose, or galactose. After 24 hours of growth, cultures were diluted and plated onto YPD agar. Resulting colonies were replica plated onto amino acid double-dropout plates tailored to the specific auxotrophic markers of the strain pair. Colonies able to grow on dropout media were scored as hybrids, while the rest were haploid parental types. This allowed for quantification of hybrid frequency in the evolving populations under each sugar condition.

## Acknowledgements

This work was funded by a DBT/Wellcome Trust (India Alliance) grant (award no. IA/S/19/2/504632) to SS. NR is supported by the Prime Minister’s Research Fellowship (PMRF ID 1301163).

